# APHIX: Analysis Pipeline for HIV-1 Isoform eXploration Using Long-read RNA Sequencing Data

**DOI:** 10.1101/2024.12.09.627634

**Authors:** Jessica L Albert, Christian M Gallardo, Bruce E Torbett

## Abstract

HIV-1 uses 4 major splice donors and 8 major splice acceptors as well as dozens of minor, cryptic, and uncharacterized splice sites to produce over one hundred distinct transcript isoforms from a single 9.2 kb genome. As a result, existing bioinformatic pipelines struggle to accurately analyze spliced HIV sequences due to the complex nature of HIV alternative splicing compared to human mRNA splicing. Previous approaches to identify HIV isoforms from long-read sequencing data used pipelines that are not publicly available, are convoluted to operate, or are locked into a specific HIV strain, which limits their wide adoption to other experimental designs or systems. To address this gap, we have developed a bioinformatic pipeline called *APHIX* that fully automates spliced isoform assignment, splice site usage quantification, and non-coding exon detection. APHIX takes a FASTQ/A of long-read transcripts and a HIV genome reference sequence and fully automates HIV isoform analysis. APHIX calculates splice site usage counts and percentages for each donor and acceptor site and their pairwise combinations, accurately assigns isoforms, and automatically identifies transcripts containing non-coding exons. APHIX is compatible with long-reads sequences generated from multiple platforms and library preps, including direct DNA and RNA sequencing. APHIX can also be adapted to multiple HIV-1 clades and strains by providing the appropriate reference sequence during bioinformatic processing. Overall, APHIX enables comprehensive processing of spliced sequences with reproducible results in a manner that is faster and easier to run compared to other methods.

## 1. Introduction

The Human Immunodeficiency Virus 1 (HIV) utilizes a complex system of alternative splicing to produce over one hundred distinct transcript isoforms from a single 9.2 kb genome^1^. To enable this range of isoform diversity, HIV has 4 major splice donors and 8 major splice acceptors as well as dozens of minor, cryptic, and novel splice sites^2^. These splice sites are used in various combinations to bring the translation start site of each isoform proximal to the shared 5’ untranslated region (UTR)^1^. The specific isoform identity of a transcript is generally determined using the ribosomal scanning principle, in which the translation start site closest to the 5’ cap determines the open-reading frame that is translated. Viral transcripts that are not spliced, referred to as unspliced (US), code for structural and enzymatic proteins (Gag and Gag-Pol) as well as acting as the viral genome^1^. Fully spliced (FS) transcripts are about 2kb in length and have all introns removed, typically consisting of Tat, Rev, and Nef transcripts^1^. There are also transcripts that are spliced but retain the donor 4 (D4) to acceptor 7 (A7) intron and are about 4kb in length. These are referred to as partially spliced (PS), and typically consist of Vif, Vpr, and Env/Vpu transcripts^1^. While most spliced isoforms are within their stated size class, there are small subsets in another spliced size class, such as the FS Vif and Vpr transcripts and PS Tat transcripts^3^.

In addition, spliced HIV transcripts can contain two small non-coding exons (NCE) that span from A1 to D2 (NCE2) and A2 to D3 (NCE3)^3^ and are sometimes included (individually or in combination) in most spliced isoforms (except in Vif and some Vpr transcripts). The purpose of these NCEs in HIV spliced transcripts has not been determined, but studies have suggested NCEs could be structured in the form of a stem loop which could have functional effects on stability, processing, or translation rate of the underlying transcript^4^. Previous studies removing the NCEs while preserving Vif and Vpr production have had conflicting results. There is some evidence that including the NCEs can increase the stability of the transcripts^5^, but other studies have found little to no difference in transcript stability as a result of NCE inclusion^6, 7^.

Given the complexity of HIV splicing, exons are reused in various combinations to create dozens of distinct isoforms, which limits the ability to unambiguously determine isoform identity. If short read sequencing is used, the resulting reads would map to multiple isoforms due to the identical 5’ and 3’ UTRs and use of the same splice sites in distinct isoforms. For example, a short read that spans the D4-A7 junction, only determines that it is from a FS transcript, but not what protein it encodes. Long-read sequencing can partially ameliorate isoform assignment ambiguities. However, if multiple priming strategies are used to amplify the isoforms, then the isoform usage ratios cannot be accurately determined due to differences in primer efficiencies. On the other hand, using a single priming strategy, such as an Oligo-d(T) strategy, to capture full-length transcripts, allows for the usage ratios of different isoforms to be identified since the single priming strategy eliminates any differences in primer efficiency^8^.

Although long-read sequencing alone is informative, many publicly available tools for isoform identification (IsoQuant^9^, Flair^10^, Pinfish^11^, etc.) were built on splicing patterns typically found in eukaryotic organisms and struggle when processing reads from viral transcripts. While typical eukaryotic splicing does have alternative exons or retained introns, the presence or absence of these exons does not typically change the identity of the resulting protein^1^. For eukaryotic transcripts, exons are generally a few hundred bases, while introns are often one to two kilobases. HIV’s use of large exons, extensive intron retention, and alternative exons creating distinct products leads to the current tools struggling to accurately identify HIV isoforms without manual parsing. Previous studies using long-read sequencing have used a wide range of post hoc analyses that are poorly described, time consuming, difficult to reliably replicate and/or restricted to specific HIV strain analysis (ex: NL43, HXB2, etc).

To improve the speed and accuracy of HIV isoform identification, we developed a Snakemake pipeline that automates the processing, identification, and analysis of HIV isoforms and calculates isoform, NCE, and splice site usage from the resulting isoforms. The resulting tool, dubbed **<u>A</u>**nalysis **<u>P</u>**ipeline for **<u>H</u>**IV **<u>I</u>**soform e**<u>X</u>**ploration (APHIX)^12^, was built on previous HIV isoform analysis methods from Gallardo et al 2022^13^ with the addition of python-based tool, *HIV_isoform_checker*^*14*^, to replace manual parsing steps. In this study, we establish the utility of APHIX in the rapid processing of long-read direct RNA and direct cDNA sequencing from cellular RNA from multiple cell lines infected with a variety of HIV strains. In addition, we demonstrate and validate that APHIX can reproduce isoform and splice site results from published data. We show that APHIX automatically identifies NCE presence in transcripts allowing for in depth analysis of an understudied aspect of HIV splicing. APHIX is publicly available via GitHub and can be set up in minutes using a prebuilt conda environment. To run the pipeline, file inputs in can be invoked in the pipeline’s configuration file and executed with a single command in the Command Line Interface (CLI). APHIX is fast and simple to use allowing for quick, reproducible isoform, splicing and NCE analysis for most HIV-1 strains with minimal adjustments.

## 2. Methods

### 2.1 APHIX pipeline

A Snakemake pipeline combining multiple programs was created to automate the analysis of isoform dynamics and splice site usages for spliced HIV isoforms from long-read sequences. The pipeline, associated manual, and test data is available at (https://github.com/JessicaA2019/APHIX). It can be initialized in minutes using a prebuilt conda environment and run in seconds using one simple command. APHIX has been extensively tested with Ubuntu 20.04.6 LTS and is currently compatible with Linux platforms with x86-64 architectures.

To run APHIX, simply adjust the variables in the config file (see Box 1) and run the following command: “snakemake --cores 4 --configfile config.yml”. The run time for APHIX is relatively quick, ranging from about 30 seconds for MinION scale dataset and about five minutes for PromethION scale datasets tested (see Supp. Table 1). For larger or more complex datasets, the number of threads/cores used by APHIX can be increased to speed up processing.

APHIX is split into 4 main sections: (1) mapping transcripts with minimap2; (2) clustering and polishing with Oxford Nanopore Technologies’ Pinfish tools; (3) preliminary isoform identification with GFFcompare; and (4) correcting isoforms with HIV_Isoform_Checker.

#### 2.1.1 Mapping Transcripts

Input fast(a/q) for the sample are mapped using minimap2^15^ preset with *-ax, splice*, and *-- secondary=no* parameters. These parameters can be altered in the “Optional parameters” section in the Snakefile. The mapped transcripts are extracted using samtools view with *-bS -F4 -h* parameters and sorted with samtools sort. The resulting bam file is then indexed with samtools index. These commands are shown in Box 2 Step 2.

#### 2.1.2 Clustering and Polishing with Pinfish^10^

The indexed bam file is then converted to a gff file using spliced_bam2gff^16^ (see Box 2 Step 3). Gff is clustered using *cluster_gff*^11^ preset with *-d 5 -p 0* parameters and the cluster size (*-c*) parameter indicated in the config file (see Box 2 Step 4). The preset parameters can be altered in the “Optional parameters” section in the Snakefile. The resulting tsv file is then polished using *polish_clusters*^11^ with the same cluster size (*-c*) parameter indicated in the config file (see Box 2 Step 5). If FASTA files were given as input, an additional *-f* parameter is automatically added to the polishing command.

#### 2.1.3 Preliminary Isoform Identification with GFFcompare

The FASTA file output by the polish step is then remapped to the reference genome (see Box 2 Step 6) and converted to a gff file (see Box 2 Step 7). To create a gff of HIV isoforms for the given reference, the input genome FASTA is mapped against NL43 using auxiliary python script *CLI_gff_converter.py* (Box 2 Step 8) which calls *Reference_aligner.py* which uses mappy, a python wrapper for minimap2^15^, to create a CIGAR string. This CIGAR string is then used to shift the NL43-based positions in the *auxillary.gff* file to be accurate to the provided reference genome. The created reference and polished gff are then cross referenced using *gffcompare*^17^ with preset *-V -M -T* parameters to attain preliminary isoform identifications (see Box 2 Step 9). The resulting annotated gtf file is then fed into *HIV_Isoform_Checker.py*^14^ with *-n True* as a preset parameter (see Box 2 Step 10). This variable and the others discussed in Box 3 can be altered in the “Optional parameters” section in the Snakefile.

#### 2.1.4 HIV_Isoform_Checker.py

*HIV_Isoform_Checker.py* uses accepted NL43 exon boundaries (EBs) and coding sequences (CDS) to confirm isoform identities. To convert the EBs and CDS to align with the given reference genome, *Reference_aligner.py* is used to create a CIGAR string which is used to shift the NL43-based positions. Then, the transcript ID, cluster size, isoform ID, starting and ending position, GFFcompare assigned class code, and EBs are pulled from the polished gtf file. This information is used to calculate coverage length, size normalized counts, and identify the presence of NCEs (if desired).

These clusters are then filtered as follows. First, any cluster with a GFFcompare assigned class code other than “=“ (exact intron chain match), “m” (retained introns), or “j” (multi-exon with at least on junction match) are eliminated. Next, any clusters that do not go beyond the full-length cutoff values (*-a* and *-z* in Box 3) are eliminated. Finally, any clusters that have gaps that are not associated with a known introns are either eliminated or filled in as a sequencing error (*-g* in Box 3).

After filtering out short or abnormal clusters, the isoform identity assigned by GFFcompare is validated by confirming the presence of the full CDS region associated with the identity. If the associated full CDS is not present, the transcript is checked against all possible full CDS regions from the 5’ end. If the transcript is shorter than the fully spliced cutoff (*-l* in Box 3), then it is checked for a full CDS as follows: tat, rev then nef. If the transcript is longer than the fully spliced cutoff, then it is checked for a full CDS as follows: vif, vpr, PS tat, then env/vpu. The first confirmed full CDS region is assigned as the corrected isoform identity.

Using the finalized isoform identifications and corrected EBs, the script counts the number of transcripts for each isoform, the number of times each individual splice acceptor/donor was used, and the number of times each acceptor-donor pair was used. These counts are used to calculate the percent usage across the sample. Outputs of HIV_Isoform_Checker and the earlier step of APHIX are summarized in Supplementary Box 1.

### 2.2 Analysis of Published data

#### 2.2.1 Analysis of Gallardo et al 2022^13^ Data with APHIX

Raw data corresponding to 10231-SC-TP-JL, 10232-SC-TP-JL, 1211-SC-TP-JL and 1212-SC-TP-JL from PRJNA801353 (NCBI Sequence Read Archive) was demultiplexed using Dorado v 0.8.2 with the *<u>dna_r9</u>*. *<u>4</u>*.*<u>1_e8_sup@v3</u>*.*<u>6</u>* model^18^. Briefly, this data is from J-lat 10.6 cells, a Jurkat-derived cell line that is latently infected with HIV, were activated with TNF-alpha for 24 hours^13^. Total cellular RNA was extracted and prepared for direct cDNA sequencing by reverse transcription, and second strand synthesis. The materials were then prepped for sequencing using ONT EXP-NBD104 and SQK-LSK109 protocols and sequenced on a MinION device as described^13^. After demultiplexing, the resulting FASTQ files and a reference FASTA of the J-Lat 10.6 HIV genome sequence from <u>MN989412.1</u> were then run through the APHIX pipeline using default threads and cluster size values. For splice donor and acceptor usage data, raw counts from APHIX were divided by the donor or acceptor usage total. Isoform usage rates were calculated by adding the transcripts for each isoform group (represented by the size of each cluster) and dividing by the sum of transcripts in the detailed APHIX output csv. These results were compared to source data for Figure 6c, 6d, and 6e from Gallardo et al 2022^13^.

#### 2.2.2 Analysis of Baek et al 2024^19^ Data with APHIX

FAST5 files corresponding to WT, Triple, A8079G, A8975C, and A8989T samples were downloaded from the European Nucleotide Archive using accession number <u>PRJEB61077</u>. Briefly, this data is from Jurkat cells that were infected with WT or mutant virus. Total cellular RNA was extracted at 96 hours and used for direct RNA sequencing using modified ONT SQK-RNA002 protocols and sequenced on a MinION device as described^19^. The files were converted to POD5^20^ and demultiplexed using Dorado v 0.8.2 with the *rna002_70bps_hac@v3* model^18^. The resulting FASTQ files and a reference FASTA of the NL43 genome sequence from <u>AF324493.2</u> were then run through the APHIX pipeline using default threads and cluster size values. For splice donor and acceptor usage data, raw counts from APHIX were summed across the replicates for each sample group and divided by the donor or acceptor usage total. Spliced isoform usage rates were calculated by adding the transcripts for each isoform group (represented by the size of each cluster) and dividing by the sum of transcripts in the detailed APHIX output csv. To calculate the unspliced transcripts, GFFcompare was run as in Box 2 Step 9 except without the -M argument so that monoexonic transcripts are allowed. The resulting gff file was then manually processed to count all the transcripts identified as “Pol” by GFFcompare that were over 8kb in length to match with Baek et al’s analysis. These results were compared to source data provided by Baek et al for figure 4d(v) and 4e^19^. To calculate the NCE usage, transcripts were binned by the presence or absence of each NCE. Then, transcript counts were separately summed for each sample group, both as a whole and within each isoform type, and divided by the sum of all transcripts.

#### 2.2.3 Analysis of direct cDNA sequencing of infected primary cells with APHIX

NL43-infected donor derived primary cells were used for direct cDNA sequencing as described^8^. Briefly, total RNA was extracted 7 days post-infection. Total cellular RNA was extracted and prepared for direct cDNA sequencing by reverse transcription, and second strand synthesis. The materials was then prepped for sequencing using ONT SQK-NBD114 protocols and sequenced on a PromethION device. The resulting FASTQ files and a reference FASTA of the NL43 genome sequence from <u>AF324493.2</u> were then run through the APHIX pipeline using default threads and cluster size values. To calculate the NCE usage, transcripts were binned by the presence or absence of each NCE. Then, transcript counts were separately summed for each replicate, both as a whole and within each isoform type, and divided by the sum of all transcripts.

## 3. Results

### Improvement of the specificity and speed of HIV-1 spliced isoform analysis from long-read RNA sequencing data using APHIX

APHIX leverages a combination of publicly available tools with custom python scripts to identify HIV-1 isoforms and analyze splice site usage **(Fig 1)**. Briefly, minimap2^15^ is used to map long-read transcripts to the provided HIV reference genome. The resulting file is clustered and polished using scripts from Oxford Nanopore Technologies’ Pinfish pipeline^11^. The resulting polished clusters are preliminarily identified with GFFcompare^17^ and then validated with HIV_Isoform_Checker^14^ which also confirms the isoform identity. On a standard computer using 4 threads, APHIX runs in a few minutes **(Supp. Table 1)** and outputs isoform, NCE, and splice site usage analyses as well as all intermediate files created by the pipeline.

**Figure 1:**
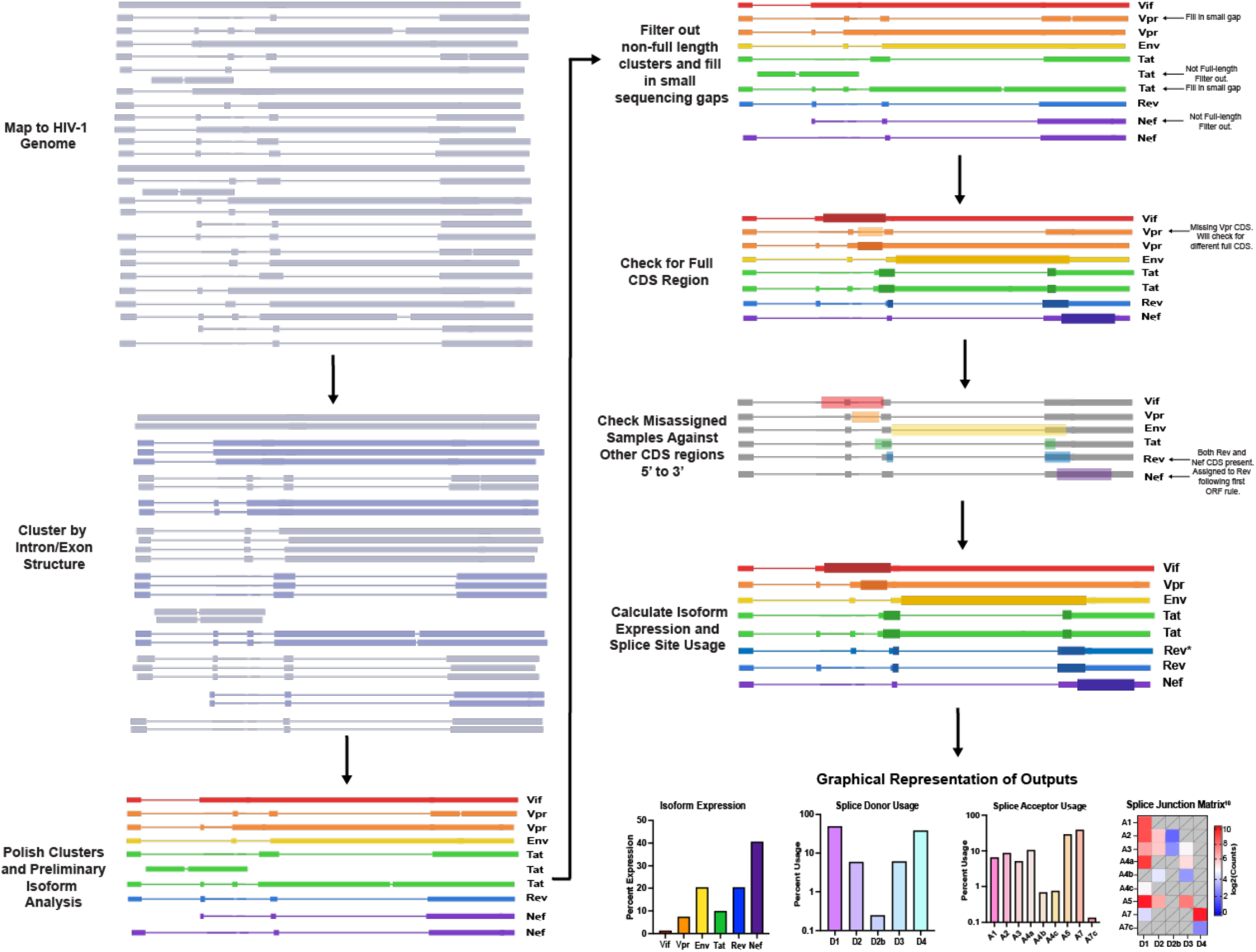
APHIX bioinformatic pipeline. Input FASTQ/A are aligned to the provided HIV reference, followed by clustering, polishing, preliminary isoform identification. These preliminary isoforms are then filtered, and their identities are confirmed resulting in the usage counts and percentages for each isoform group as well as each donor site, acceptor site, and pairwise combination.

### Validation of automated APHIX results through comparison with original non-automated results from Gallardo et al. 2022

We first determined whether APHIX can accurately assign isoform identities and calculate splice site usage in concordance to the published direct cDNA sequencing data in Gallardo et al. 2022^13^. APHIX was able to replicate isoform usage +/- 3.2% with correlations for each replicate showing R^2^ values greater than 0.99 **(Fig. 1A)**. For splice donor usage, APHIX was able to replicate the original data +/- 3.3% with correlations for each replicate with R^2^ values greater than 0.99 **(Fig. 1B)**. For splice acceptor usage, APHIX was able to replicate the original data +/- 9.0% with correlations for each replicate with R^2^ values greater than 0.97 **(Fig. 1C)**. However, the A4a and A4b usage shows inconsistencies with the original values, which we traced to a suboptimal exon boundary threshold that was inadvertently used in the Gallardo et al. 2022 manuscript. In APHIX, exon boundary thresholds were optimized so that exon boundaries that are in close proximity to each other (as few as 6 bp away) can be precisely clustered. When misassigned A4a and A4b junctions from the original manuscript are excluded, APHIX was able to replicate the original data +/- 4.5% with correlations for each replicate with R^2^ values greater than 0.99. This data provides critical validation that APHIX has an improved ability to accurately assign isoform identities and perform splice site usage calculations compared to the original pipeline.

### Evaluation of APHIX through Isoform and Splice Site Usage rates for published Direct RNA sequencing data

To evaluate whether APHIX can accurately assign isoform identities and calculate splice site usage for direct RNA sequencing (DRS) data, APHIX was used to process data from Baek et al 2024 and was compared to the processed datapoints reported in their manuscript^19^. APHIX was able to replicate isoform usage +/- 6.1% with correlations for each group with R^2^ values greater than 0.99 except for group A8975C with R^2^ values greater than 0.98 (Fig. 1A). There are minor differences between Baek et al’s isoform analysis and APHIX results which is most notable in Nef which is due to it being the most highly expressed isoform and thus having the largest difference. For splice donor usage, APHIX was able to replicate the Baek et al 2024 data +/- 1.3% with correlations for each replicate with R^2^ values greater than 0.99 (Fig. 1B). For splice acceptor usage, APHIX was able to replicate the Baek et al 2024 data +/- 2.2% with correlations for each replicate with R^2^ values greater than 0.99 (Fig. 1C). Our findings provides validation that APHIX has an improved ability to quickly assign isoform identities and perform splice site usage calculations, when compared to the Baek et al’s methods.

### Identification of NCE usage by APHIX shows preferential addition of NCEs to certain isoform classes

APHIX provides outputs that indicate the presence or absence of NCE 2, NCE 2b, and NCE 3 for each cluster during the *HIV_Isoform_Checker* step by default (see Methods section 2.1.4). This allows for rapid analyses of NCE usage across a sample and within isoform groups. APHIX analysis of direct cDNA sequencing of NL43 infected donor derived primary cells^8^ found that about 26% of transcripts contained one or more of the NCEs, with one donor, having only 22% of transcripts with a NCE (Fig. 4A). Most of the NCEs present were NCE3. Analysis of the Baek et al 2024 data found usage of any NCE in about 30% of transcripts with a majority of NCE containing reads having the NCE2 variant (Fig. 4C). The Triple mutant had the highest usage of NCEs (35%). When separated by isoform class, 2kb class isoforms had higher rates of transcripts with NCEs (34% and 40%) compared to 4kb transcripts (11% and 19%) in both the DCS (fig 4B) and DRS (Fig, 4D) data, respectively. Interestingly, Tat had a significantly higher rate of NCE inclusion (one-way ANOVA, P <0.0001 for both DCS and DRS), even compared to the other 2kb class isoforms, Nef (P=0.0004 for DCS, P<0.0001 DRS) and Rev (P<0.0001 for both DCS and DRS).

**Figure 2:**
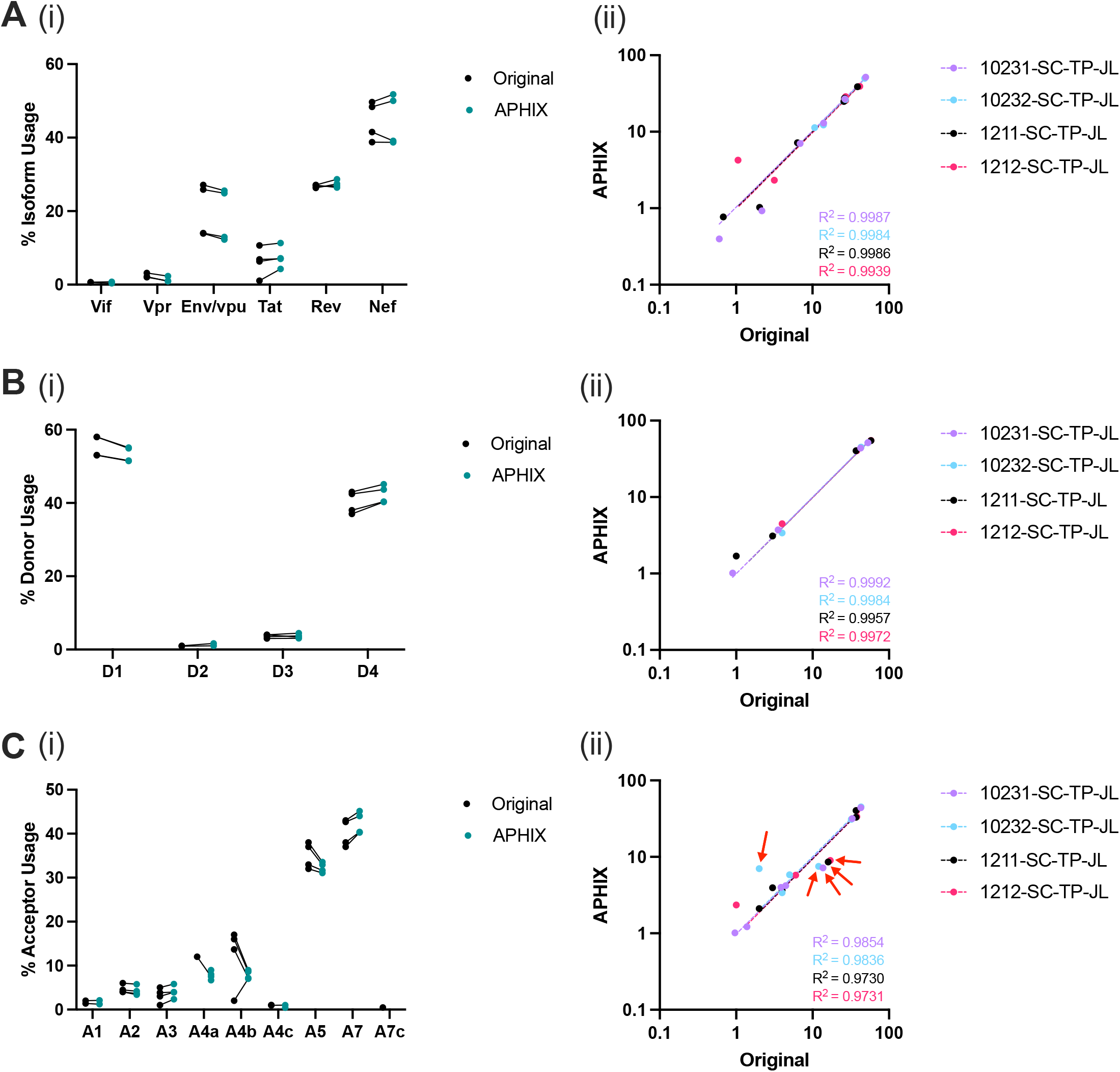
Comparison of original analysis and APHIX. **(i)** Usage fractions for the original analysis (black) and APHIX (teal) and **(ii)** correlation graph of the original analysis on the x-axis and APHIX on the y-axis with each color as a different replicate for **(A)** isoforms, **(B)** splice donors, and **(C)** splice acceptors. Red arrows panel C-ii designate clusters A4a and A4b which are processed differently between APHIX and Gallardo et al 2022.

**Figure 3:**
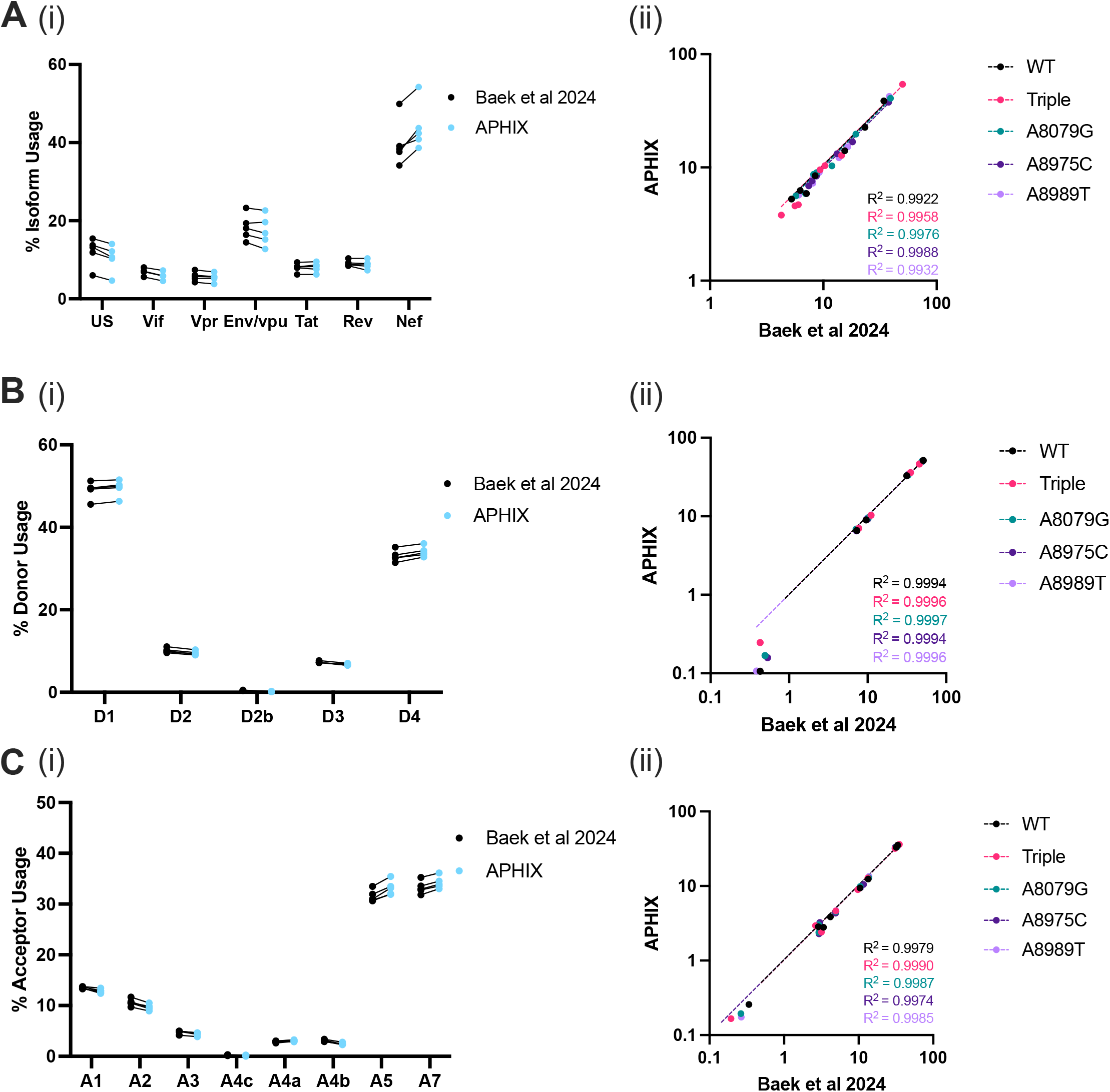
Comparison of Baek et al 2024 data and APHIX. **(i)** Usage fractions for the Baek et al 2024 data (black) and APHIX (blue) and **(ii)** correlation graph of the Baek et al 2024 data on the x-axis and APHIX on the y-axis with each color as a different sample type for **(A)** isoforms, **(B)** splice donors, and **(C)** splice acceptors.

**Figure 4:**
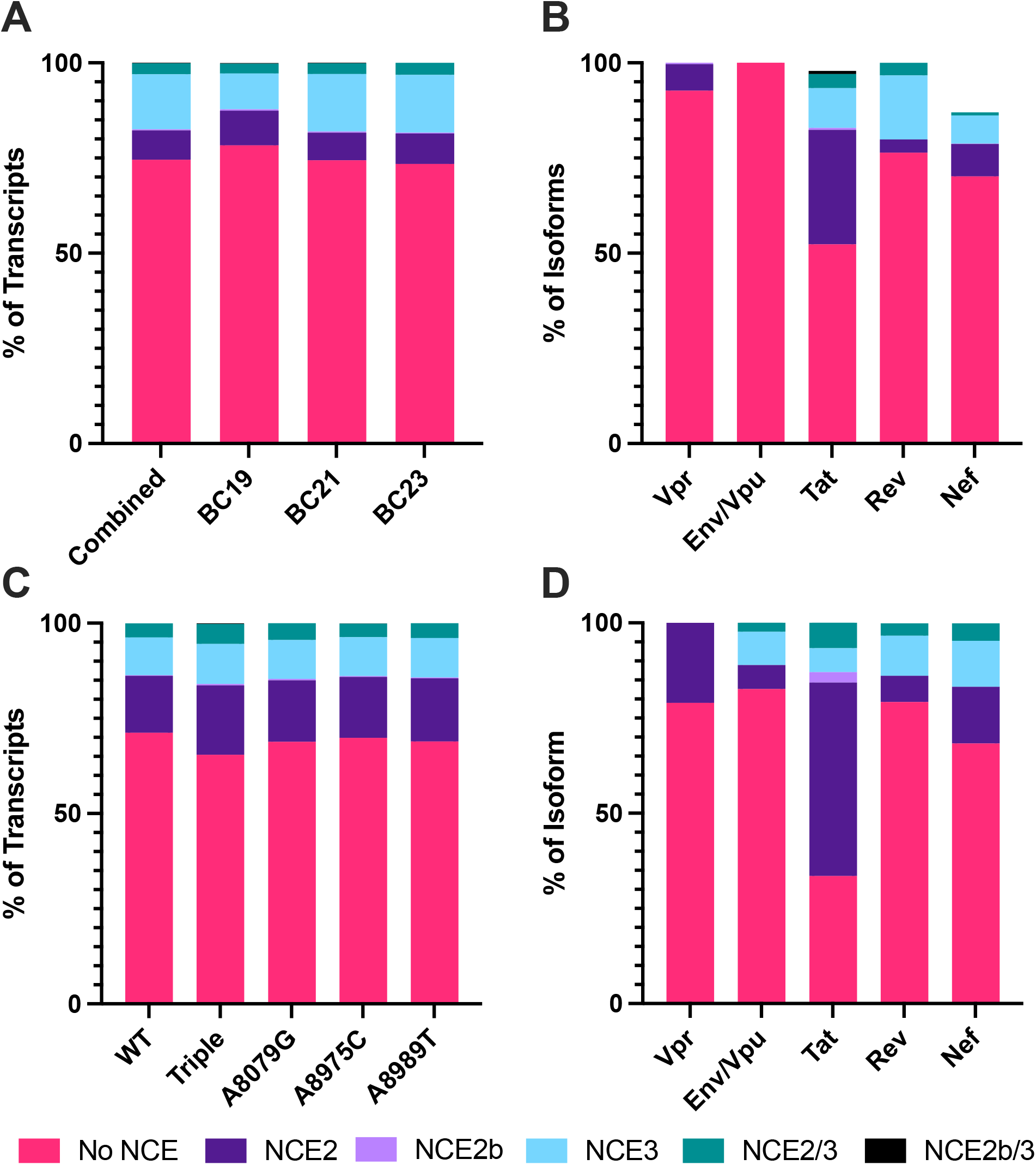
NCE usage in primary cell DCS and Baek et al 2024 data. **(A)** Rate of NCE inclusion across transcripts in primary cells. **(B)** Rate NCE inclusion for each isoform class in primary cells. **(C)** Rate of NCE inclusion for each sample group from Baek et al 2024. **(D)** Rate of NCE inclusion in the WT group for each isoform class from Baek et al 2024.

## 4 Discussion and Future Directions

In this study, we introduce and benchmark APHIX, an automated Snakemake pipeline for isoform analyses of spliced HIV-1 long-read sequences. APHIX produces extensive isoform analyses rapidly and is completely unsupervised. It is easily deployable using a prebuilt conda environment, requires minimal bioinformatic experience, and is run with a single command. APHIX is also easily modifiable though variables found in the config file. APHIX has been tested with multiple HIV strains, including those with large insertions and deletions within the HIV genome. APHIX can use a multitude of long-read sequencing methodologies and library preparations, including direct cDNA and RNA preparations shown above and amplification-based methods^8^. While APHIX has been built and tested using ONT sequencing methodologies, sequences from other long-read platforms such as PacBio, should be compatible with APHIX.

We have shown our approach determines isoform, splice site, and NCE usage in direct RNA and cDNA sequencing data from multiple cell lines and HIV strains. This allows improved speed and reproducibility of spliced HIV isoform analyses. Moreover, the pipeline is more accurate than the original non-automated analysis due to the increased stringency of isoform clustering^8^. This allows for differentiation of splice donors and acceptors that are within close proximity of each other, such as A4a and A4b which are six base pairs apart. There were some inconsistencies in isoform percentages between APHIX and Baek et al 2024. These differences are likely due to the increased stringency of APHIX, which requires at least three transcripts per intron/exon pattern and a limited preset list of well-described splice donors and acceptors. This was seen most notably in quantification of Nef, which, being the most prevalent isoform, leads to the differences in quantification being more evident. Although there are differences seen between analysis pipelines, APHIX maintains the differences in isoform and splice site usage observed between WT and the mutants found in Baek et al 2024.

APHIX automatically provides identification of NCEs within isoforms which allowed identification of trends in NCE usage, an advantage of our pipeline not found in existing pipelines or manual parsing methods. Most notably, APHIX analysis found the increased usage of NCEs in FS transcript class, particularly in Tat. Both data sets used NL4-3 virus, so a further exploration of this phenomenon in a more diverse array of cell lines and HIV strains is warranted. However, if this trend holds in other strains, one possible explanation of the NCEs preferentially being found in 2kb transcripts is that by splicing out the D4-A7 intron, there is a shift in the transcript structure that makes the NCEs more accessible for splicing, possibly through shifts in splice donor accessibility or displacement of splicing repressors. Given that Tat is closest to these NCEs, there may be a greater effect on the accessibility of the associated splice sites. Another possibility is that the reduction in splicing repression needed to splice to A3^21^, which is used in Tat and is the least utilized acceptor^3^, also reduces the splicing inhibition on A1 and A2, which are used to create NCE2 and NCE3 respectively. More research exploring how RNA structure and splice repressor binding shifts during the splicing process would be needed to determine the mechanism of NCE inclusion preferences.

The ability of APHIX to take a FASTQ of long-read HIV sequences and quickly perform a multitude of isoform analyses can help to simplify the discovery of complex patterns related to HIV RNA processing. APHIX currently only processes spliced HIV reads, but future iterations of the pipeline will include the ability to quantify unspliced^2^ and antisense transcripts^22^. Along with the expanded isoform classes, an option to concurrently quantify isoform productivity (i.e. the ability of a transcript to code for a translatable protein) for the transcripts that pass the previous filters will be implemented. APHIX will also be expanded to process a wider range of retroviruses, including HIV-2, SIV, HTLV-1, MMLV, and simian foamy virus. APHIX is currently only available for Linux x86-64 operating systems, but future versions will be compatible with ARM64 architectures and Windows operating systems. Additionally, a batch mode to run multiple samples at once is planned to even further improve the speed and ease of use of APHIX.

## Funding

National Institute of Allergy and Infectious Diseases [U54AI170855 to B.E.T]; Washington Research Foundation [P1-0376 to C.M.G].

## Figures and boxes

### Box 1

Config File Parameters

A brief description of the parameters set in the config file before running APHIX. Variables shown are for provided test data. The variables to adjust are bolded.

1. Name of output folder: Designates the prefix of the output folder that will be made in the working directory and the name that the output files in said folder will be given. Use “-” or “_” instead of spaces and do not add any periods.sample_name: **APHIX_test**
2. Input FASTQ file: Designates the input location of the FASTQ or FASTA file to be processed. input_fastq: **test_files/APHIX_test.fastq**
3. Reference genome: Designates the input location of the FASTA file of the reference genome to be used for processing. Do not remove the quotation marks.reference_fasta: “**test_files/NL43.fa**”
4. Name for reference gff to be created: Designates the name of the gff file created for the above reference genome to be used by gffcompare. This file will at “APHIX_test_isoform_analysis/7_gff_compare/”. Do not remove the quotation marks.reference_gff_name: “**NL43.gff**”
5. Threads: Designates the number of threads used by multiple steps in the APHIX pipeline. This value should match the “--cores” argument in the Snakemake command. If “--cores” and threads do not match, the program will cap out at the lower of the two values. Default is 4. threads: **4**
6. Is the input file a FASTQ file?: Designates if the input file given above is a FASTQ or a FASTA file. If the file is a FASTQ, variable should be “True”. If the file is a FASTA, variable should be “False”. Default is True. input_is_fastq: **True**
7. Cluster Size: Designates the cluster size given to cluster_gff and polish_clusters in APHIX pipeline. A higher the cluster size leads to the isoforms output by APHIX having higher specificity, at the expense of lowering sensitivity. Default is 3.

cluster_size: **3**

### Box 2

*Summary of commands run by Snakemake pipeline*

A brief description of rules and associated commands performed by the APHIX Snakemake pipeline to process test data.

1. ref_no_header: Removing any headers from the input genome reference FASTA to prevent downstream errors. seqkit seq -i -w 0 test_files/NL43.fa > APHIX_test_ref_no_header.fa
2. map_fq: Mapping input fast(q/a) file to the provided genome reference using the “splice” setting with no secondary alignments. Reads that map to the reference (-F4) are then sorted and indexed. -t and -@ refer to the number of threads used. minimap2 -ax splice -t 4 --secondary=no APHIX_test_ref_no_header.fa test_files/APHIX_test.fastq | samtools view -bS -F4 -h -@ 4 - | samtools sort -@ 4 - > APHIX_test.bam && samtools index -@ 4 APHIX_test.bam
3. bam2gff: Converts the bam file produced above into a gff file for clustering. spliced_bam2gff -M APHIX_test.bam > APHIX_test.gff
4. cluster: Reads are then clustered based on exon/intron pattern with a 5 base exon boundary tolerance (-d). Only clusters with at least 3 reads (-c) will be output. -c is determined by the cluster size given in the config.yml file. cluster_gff -c 3 -d 5 -p 0 -t 4 -a APHIX_test_clusters.tsv APHIX_test.gff > APHIX_test_clusters.gff
5. polish: Clusters are then polished into consensus exon/intron boundaries. polish_clusters -a APHIX_test_clusters.tsv -c 3 -t 4 -o APHIX_test_polished.fa APHIX_test.bam
6. map_clusters: Polished clusters are then re-mapped to the genome reference as above. minimap2 -ax splice -t 4 --secondary=no APHIX_test_ref_no_header.fa APHIX_test_polished.fa | samtools view -bS -F4 -h -@ 4 - | samtools sort -@ 4 - > APHIX_test_polished.bam && samtools index -@ 4 APHIX_test_polished.bam
7. bam2gff_clusters: Converts the bam file produced above into a gff file for clustering. spliced_bam2gff -M APHIX_test_polished.bam > APHIX_test_polished.gff
8. create_gff_ref: A gff file for the genome reference is created by mapping the given reference against the NL43 sequence and converting the base pair positions/boundaries to match the given reference. python3 auxillary_scripts/CLI_gff_converter.py auxillary_scripts/auxillary.gff NL43.gff APHIX_test_ref_no_header.fa
9. gff_compare: The polished gff is compared against the created reference gff to assign preliminary isoform identities and remove monoexonic reads (-M). gffcompare -V -M -T -r NL43.gff APHIX_test_polished.gff -j gffcmp_novel_junction.tab
10. HIV_Isoform_Checker.py: The preliminarily annotated gff file is then filtered to remove non-full-length clusters and non-canonical gaps over 15 base pairs. These filtered clusters are then screened to confirm the full CDS of the predicted isoform is present. If the assigned full CDS is not present, then the cluster is check against the other isoform identities and the first full CDS from the 5’ direction is used to reassign the correct isoform identity.

HIV_Isoform_Checker -n True APHIX_test.annotated.gff checked_isoforms APHIX_test_ref_no_header.fa

### Box 3

<u>*HIV_Isoform_Checker variables*</u>

A brief description of the modifiable parameters available for HIV_Isoform_Checker. Commands with default values are listed.

1. Gap Tolerance: If a read has a gap longer than this value that is not associated with known splice sites, the read will be filtered out as a major sequencing error. Any gaps smaller than or equal to this value will be filled in as minor sequencing error for isoform confirmation purposes. -g 15 or --gap_value 15
2. Starting Base Pair Position: Sets maximum starting bp in order to ensure the presence of the first exon and by proxy a full length read. If the read does not have any bases at or before this value, then it is filtered out as a not full-length read. -a 700 or --startBP 700
3. Ending Base Pair Position: Sets minimum ending bp in order to ensure the presence of the last exon and by proxy a full length read. If the read does not have any bases at or after this value, then it is filtered out as a not full-length read. -z 9500 or --endBP 9500
4. Fully Spliced Cutoff: Sets maximum fully spliced transcript length. If a read is longer than this value, then it will be considered partial spliced. This is based off HIV-1 references without any major insertions or deletions. If your sequences have larger deletions (ex: deltaENV or deltaMA) or insertions (ex: multiple markers such as eGFP), this value may need to be adjusted to be shorter or longer, respectively. -l 2500 or --lengthFS 2500
5. NCE Identification: When set to True, the output csv file will have y/n columns for the presence of NCEs. -n False or --NCE_value False

## Notes

### Competing Interest Statement

The authors have declared no competing interest.

