## Supplemental Figures for "APHIX: Analysis Pipeline for HIV-1 Isoform eXploration Using Long-read RNA Sequencing Data"

Table of APHIX Details

| Source | File | Genome | Cluster Size | # total sequences | # HIV sequences | Run Time |
|---|---|---|---|---|---|---|
| Baek | WT_rep1.fastq | NL43_Complete.fa | 3 | 163,748 | 1,219 | 5s |
| Baek | WT_rep2.fastq | NL43_Complete.fa | 3 | 951,155 | 6,227 | 17s |
| Baek | WT_rep3.fastq | NL43_Complete.fa | 3 | 1,417,887 | 7,594 | 25s |
| Baek | WT_rep4.fastq | NL43_Complete.fa | 3 | 594,978 | 3,678 | 11s |
| Baek | Triple_rep1.fastq | NL43_Complete.fa | 3 | 1,185,407 | 8,883 | 22s |
| Baek | Triple_rep2.fastq | NL43_Complete.fa | 3 | 1,600,114 | 8,848 | 29s |
| Baek | Triple_rep3.fastq | NL43_Complete.fa | 3 | 1,038,811 | 5,821 | 18s |
| Baek | A8079G_rep1.fastq | NL43_Complete.fa | 3 | 1,159,945 | 5,916 | 19s |
| Baek | A8079G_rep2.fastq | NL43_Complete.fa | 3 | 995,031 | 4,726 | 16s |
| Baek | A8079G_rep3.fastq | NL43_Complete.fa | 3 | 691,272 | 2,293 | 10s |
| Baek | A8975C_rep1.fastq | NL43_Complete.fa | 3 | 342,984 | 1,911 | 8s |
| Baek | A8975C_rep2.fastq | NL43_Complete.fa | 3 | 999,284 | 5,015 | 18s |
| Baek | A8975C_rep3.fastq | NL43_Complete.fa | 3 | 987,187 | 4,756 | 16s |
| Baek | A8989T_rep1.fastq | NL43_Complete.fa | 3 | 716,450 | 5,306 | 16s |
| Baek | A8989T_rep2.fastq | NL43_Complete.fa | 3 | 992,008 | 5,363 | 18s |
| Baek | A8989T_rep3.fastq | NL43_Complete.fa | 3 | 1,053,905 | 4,618 | 15s |
| Gallardo | 10231-SC-TP-JL.fastq | R7_validated.fa | 3 | 510,766 | 4,929 | 12s |
| Gallardo | 10232-SC-TP-JL.fastq | R7_validated.fa | 3 | 207,244 | 2,020 | 8s |
| Gallardo | 1211-SC-TP-JL.fastq | R7_validated.fa | 3 | 494,631 | 6,440 | 12s |
| Gallardo | 1212-SC-TP-JL.fastq | R7_validated.fa | 3 | 365,217 | 4,112 | 9s |
| Gallardo | Barcode19.fastq | NL43_Complete.fa | 3 | ---- | 28,618 | 46s |
| Gallardo | Barcode21.fastq | NL43_Complete.fa | 3 | ---- | 113,782 | 4m48s |
| Gallardo | Barcode23.fastq | NL43_Complete.fa | 3 | ---- | 69,626 | 2m18s |

Table of CDS Coordinates

| Isoform Name | NL43 Coordinates |
|---|---|
| Gag | 790-2289 |
| Pol | 2085-5093 |
| Vif | 5041-5616 |
| Vpr | 5559- 5846 |
| Tat | 5830-6044, 8369-8411 |
| Env | 6221-8782 |
| Rev | 5969-6044, 8369-8640 |
| Nef | 8787-9404 |

Table of Splice Donors and Acceptors

| Name | NL43 Coordinate |
|---|---|
| D1 | 743 |
| D2 | 4962 |
| D2b | 5058 |
| D3 | 5463 |
| D4 | 6044 |
| A1 | 4913 |
| A2 | 5390 |
| A3 | 5777 |
| A4a | 5954 |
| A4b | 5960 |
| A4c | 5936 |
| A5 | 5976 |
| A7 | 8369 |
| A7c | 8345 |

Table of Manual vs APHIX % Usage Data

| | 10231-SC-TP-JL | | 10232-SC-TP-JL | | 1211-SC-TP-JL | | 1212-SC-TP-JL | |
|---|---|---|---|---|---|---|---|---|
| | Manual | APHIX | Manual | APHIX | Manual | APHIX | Manual | APHIX |
| Vif | 0.605 | 0.397 | 0.000 | 0.000 | 0.680 | 0.769 | 0.000 | 0.000 |
| Vpr | 2.177 | 0.927 | 0.000 | 0.000 | 2.040 | 1.026 | 3.170 | 2.326 |
| Env | 14.027 | 12.980 | 13.930 | 12.264 | 25.850 | 24.872 | 27.110 | 25.581 |
| Tat | 6.892 | 7.020 | 10.660 | 11.321 | 6.350 | 7.179 | 1.060 | 4.264 |
| Rev | 26.602 | 26.887 | 27.050 | 26.415 | 26.300 | 27.436 | 27.110 | 28.682 |
| Nef | 49.698 | 51.788 | 48.360 | 50.000 | 38.780 | 38.718 | 41.550 | 39.147 |
| D1 | 53.065 | 51.558 | 53.000 | 51.456 | 58.000 | 54.930 | 58.000 | 55.128 |
| D2 | 0.901 | 1.016 | 0.000 | 0.000 | 1.000 | 1.690 | 0.000 | 0.000 |
| D3 | 3.546 | 3.726 | 4.000 | 3.398 | 3.000 | 3.099 | 4.000 | 4.487 |
| D4 | 42.488 | 43.699 | 43.000 | 45.146 | 37.000 | 40.282 | 38.000 | 40.385 |
| A1 | 1.386 | 1.220 | 0.000 | 0.000 | 2.000 | 2.113 | 0.000 | 0.000 |
| A2 | 4.461 | 4.201 | 4.000 | 3.398 | 4.000 | 3.662 | 6.000 | 5.769 |
| A3 | 3.858 | 3.997 | 5.000 | 5.825 | 3.000 | 3.944 | 1.000 | 2.350 |
| A4a | 0.000 | 6.707 | 12.000 | 7.524 | 0.000 | 8.028 | 0.000 | 8.974 |
| A4b | 13.683 | 7.182 | 2.000 | 7.039 | 16.000 | 8.592 | 17.000 | 8.974 |
| A4c | 0.964 | 1.016 | 1.000 | 0.000 | 0.000 | 0.423 | 1.000 | 0.000 |
| A5 | 32.972 | 31.640 | 32.000 | 31.068 | 37.000 | 32.958 | 38.000 | 33.547 |
| A7 | 42.676 | 44.038 | 43.000 | 45.146 | 37.000 | 40.282 | 38.000 | 40.385 |
| A7c | 0.000 | 0.000 | 0.000 | 0.000 | 0.476 | 0.000 | 0.000 | 0.000 |

Table of Baek vs APHIX % Usage Data

| | WT | | Triple | | A8079G | | A8975C | | A8989T | |
|---|---|---|---|---|---|---|---|---|---|---|
| | Baek | APHIX | Baek | APHIX | Baek | APHIX | Baek | APHIX | Baek | APHIX |
| Env | 21.939 | 22.625 | 13.679 | 13.679 | 18.209 | 19.652 | 17.063 | 16.831 | 15.474 | 15.199 |
| Nef | 32.206 | 38.632 | 47.206 | 47.206 | 36.803 | 40.848 | 35.528 | 43.733 | 36.087 | 42.401 |
| US | 14.566 | 14.051 | 5.685 | 5.685 | 11.183 | 10.331 | 12.476 | 10.685 | 12.941 | 12.169 |
| Rev | 7.984 | 7.332 | 9.785 | 9.785 | 8.203 | 8.998 | 8.199 | 8.384 | 8.682 | 9.019 |
| Tat | 5.898 | 6.241 | 8.853 | 8.853 | 7.706 | 8.683 | 7.510 | 7.594 | 7.709 | 8.264 |
| Vif | 6.689 | 5.871 | 5.298 | 5.298 | 6.548 | 5.874 | 6.630 | 5.895 | 7.599 | 7.254 |
| Vpr | 4.927 | 5.249 | 4.028 | 4.028 | 5.389 | 5.614 | 7.010 | 6.879 | 5.709 | 5.696 |
| D1 | 51.234 | 51.529 | 45.584 | 45.584 | 49.495 | 49.974 | 49.535 | 50.299 | 49.286 | 49.659 |
| D2 | 9.589 | 9.010 | 11.075 | 11.075 | 10.283 | 9.627 | 10.055 | 9.185 | 9.796 | 9.352 |
| D2b | 0.427 | 0.106 | 0.428 | 0.428 | 0.493 | 0.168 | 0.532 | 0.158 | 0.386 | 0.107 |
| D3 | 7.275 | 6.572 | 7.701 | 7.701 | 7.143 | 6.858 | 7.279 | 6.551 | 7.222 | 6.520 |
| D4 | 31.474 | 32.783 | 35.212 | 35.212 | 32.586 | 33.372 | 32.600 | 33.808 | 33.311 | 34.362 |
| A1 | 13.439 | 12.417 | 13.293 | 13.293 | 13.675 | 12.927 | 13.410 | 12.557 | 13.747 | 13.432 |
| A2 | 10.464 | 9.388 | 9.721 | 9.721 | 10.523 | 9.873 | 11.659 | 10.502 | 10.740 | 9.583 |
| A3 | 4.188 | 3.846 | 4.948 | 4.948 | 4.970 | 4.529 | 4.918 | 4.338 | 4.960 | 4.651 |
| A4c | 0.338 | 0.257 | 0.194 | 0.194 | 0.265 | 0.194 | 0.145 | 0.000 | 0.266 | 0.174 |
| A4a | 2.882 | 2.847 | 2.634 | 2.634 | 2.909 | 3.028 | 3.009 | 3.249 | 3.029 | 3.221 |
| A4b | 3.371 | 2.786 | 3.172 | 3.172 | 2.920 | 2.394 | 2.937 | 2.283 | 3.084 | 2.379 |
| A5 | 33.481 | 35.433 | 30.796 | 30.796 | 31.934 | 33.489 | 30.985 | 33.158 | 30.634 | 32.037 |
| A7 | 31.836 | 33.025 | 35.243 | 35.243 | 32.804 | 33.566 | 32.938 | 33.913 | 33.540 | 34.523 |

Supplementary Box 1: File Output structure

A map of the file structure and a brief description intermediate and output files of APHIX using test data is given.

APHIX_test_isoform_analysis

- 1_mapped_reads
    - APHIX_test.bam
        - Bam file output of mapping from step 2 of Box 2.
    - APHIX_test.bam.bai
        - Index file of APHIX_test.bam.
    - APHIX_test_ref_no_header.fa
        - Fasta of input genome reference with any header removed from step 1 of Box 2.
- 2_GFF
    - APHIX_test.gff
        - GFF file output of bam to GFF conversion from step 3 of Box 2.
- 3_clustering
    - APHIX_test_clusters.gff
        - GFF file output of clustering from step 4 of Box 2.
    - APHIX_test_clusters.tsv
        - TSV file output of clustering from step 4 of Box 2.
- 4_polishing
    - APHIX_test_polished.fa
        - Fasta file output of all polished clusters from step 5 of Box 2.
- 5_polish_map
    - APHIX_test_polished.bam
        - Bam file output of mapping from step 6 of Box 2.
    - APHIX_test_polished.bam.bai
        - Index file of APHIX_test_polished.bam.
- 6_polish_gff
    - APHIX_test_polished.gff
        - GFF file output of bam to GFF conversion from step 7 of Box 2.
- 7_gff_compare
    - APHIX_test.annotated.gtf
        - GFF file output of gffcompare from step 9 of Box 2.
    - APHIX_test.loci
        - A file of read identities to each super locus. Since HIV only has one super locus, this file is accessory.
    - APHIX_test_novel_junction.tab
        - A tab separated file containing all novel splice junctions not present in the reference gff with their associated cluster ids.
    - APHIX_test.stats
        - File containing statistics related to the accuracy, sensitivity, and precision of input compared to the reference.
    - APHIX_test.tracking
        - Compares transcripts across input samples. Since APHIX only takes one input file, this file is accessory.
    - NL43.gff
        - GFF file output of step 8 of Box 2.
- 8_corrected_isoforms
    - APHIX_test_altered.txt
        - A list of all sample IDs with altered exons to fill in small gaps.

APHIX_test.csv

- A csv containing the sample ID, cluster size, isoform ID, starting and ending base pair, gffcompare class code, exon boundaries, coverage length, size normalized counts, presence of NCEs (optional), and any additional notes for each sample.

APHIX_test_fail.txt

- A list of all sample IDs that failed one or more filters.

APHIX_test_isoform_counts.csv

- A csv of the usage counts and percentages for each isoform type.

APHIX_test.log

- A log of the arguments given to the script.

APHIX_test_pass.txt

- A list of all sample IDs that passed all filters.

APHIX_test_ref_coordinates.txt

- A list of splice site and CDS coordinates for given reference.

APHIX_test_splice_site_usage.csv

- A csv of the usage counts and percentages for each donor site, acceptor site, and pairwise splice site combination.
